# Aridity and coexistence with vascular plants determine the dynamics of coastal dune bryophyte communities

**DOI:** 10.1101/2024.10.29.620786

**Authors:** Z. Varela, S. Chozas, S. Lobo Dias, P. Giráldez, J. Monteiro, C. Branquinho, J. Hortal

## Abstract

The bryophyte communities of Atlantic coastal dunes help stabilising these habitats by promoting nutrient fixation, contributing to soil consolidation and enhancing water retention. Because of climate change and sea level rise, negative impacts on this dune vegetation have already been recorded, including habitat loss, shifts in species distribution and the spread of invasive bryophyte species. Moreover, in this stressful environment, biotic interactions may play a key in shaping plant diversity at these scales. We characterised coastal dune vegetation in 32 sampling sites along a latitudinal aridity gradient across the western Iberian Atlantic. We aimed to assess the potential interactions among moss species and between mosses and vascular plants, and analyse their relationship with abiotic factors to explain the community dynamics of dune bryophytes, and forecast how these communities may respond to climate change. Our results showed that the moss community is mainly influenced by aridity and temperature, with biotic interactions playing a minor, yet significant, role. As aridity increases, moss cover in these dune environments will decrease, and interactions with other plants are unlikely to compensate for this decline, thus in a climate change scenario we expect a decrease in coastal bryophyte community as well as the ecosystem services they provide; simultaneously the spread of the alien moss species *Campylopus introflexus*, linked to lower aridity, may also slow down. Consequently, the changing climate will shift optimal conditions for moss species to higher latitudes, pushing competition further north

## 1. Introduction

Coastal areas, although accounting for less than 15% of the Earth’s surface (European Environment Agency, EEA 1999), are home to more than 60% of the worldwide population, including one-third of Europe’s population (EEA 1999, 2021). These areas are of great economic and ecological value for many countries as they provide many important ecosystem services for human well-being, such as nutrient cycling, food production, habitat/shelter provision, natural barriers to erosion, water quality control and breeding grounds (Airoldi et al., 2007, Ruiz-Frau et al, 2020). According to the EEA, the extent of dune areas in Western and North-Western Europe has been reduced by 40% over the last century. This reduction is mainly related to urban development, recreational use, and reforestation, which took place from the mid-1970s. Therefore, since 1992, the marine dunes of the Atlantic, North Sea and Baltic have been considered as habitats of Community interest for conservation (Directive 92/43/EEC).

Low water availability, high salinity, nutrient-poor substrates and constant movement characterise Atlantic coastal dunes. They play a critical ecological role in stabilising and protecting coastal areas (European Commission, 2015). But these natural disturbances to which they are subjected, together with anthropogenic disturbances, have a direct impact on their fragile biodiversity (they are highly specialised areas of flora and fauna) and on the sustainable exploitation of resources (Brown and McLachlan, 2010). The problem is that these disturbances may be increased by climate change (e.g. sea level rise and increased storms), increasing urban sprawl and the invasion of alien plant species (Schlacher et al. 2008; Miller et al. 2010; Santoro et al. 2012). As a result, coastal dunes will become much more threatened, even to the point of disappearing (Gómez-Pina et al., 2002), causing incalculable ecological and economic losses in these areas.

Vegetation plays an important role in the formation, functioning and stability of coastal dunes, as its interaction with wind is a key process for dune development and its dynamics (Ranwell, 1972; Carter, 1995). Among dune vegetation, bryophytes are an important part as they can activate nutrient fixation processes, stabilise the dune surface, contribute to soil consolidation and help retain water (Murru et al., 2018). Indeed, the ecological importance of bryophytes in the structure and dynamics of Atlantic coastal dune vegetation has already been highlighted in several studies over the years (e.g. Robbins 1953-1954; Bonnot 1971; Magnusson 1983; Jun & Rozé 2004; Murru et al., 2018). However, as a consequence of climate change and sea level rise, negative effects on dune vegetation have already been reported, such as habitat loss, distribution changes, and the presence of invasive bryophyte species such as *Campylopus introflexus* (Rhind et al., 2001, Mendoza-González et al., 2013; Sérgio et al., 2018).

Under a stressful environment, vegetation might be subject to facilitative interactions between species. Facilitation is a key biotic interaction in shaping patterns of plant diversity at fine scales (Brooker et al. 2008; Forey et al. 2010) that occurs when one plant species enhances the germination or growth conditions of another (Forey et al. 2009). Some authors have already studied this inter-species facilitation in dune communities (e.g Shumway, 2000; Martínez, 2003; Maltez-Mouro et al., 2010; Vaz et al. 2013; Doxford et al. 2013) focusing mainly on the Stress Gradient Hypothesis (SGH; Bertness and Callaway, 1994). According to this SGH, facilitation and competition between species are considered important at opposite ends of stress gradients, although some authors have found competition even at high levels of environmental severity (e.g. Armas and Pugnaire 2009; Ariza and Tielborger 2011). Similarly, Doxford et al. (2013), observed that there can also be extreme spatiotemporal variations in the direction of interactions, and indeed showed that facilitation or competition between bryophytes and annual plants in dunes is correlated with population growth rate: at low growth, facilitation dominates, while at high population growth, competition dominates. Even if it is not the same ecosystem, other authors such as Monteiro et al., 2024 also reported that the composition of bryophytes in Pyrenean oak forests after fire stress is in balance between the competitive and colonisation abilities of moss species and the cover of vascular plants, confirming the significant role of biotic interactions in the composition of dune bryophyte communities.

Studying spatial climate gradients can help us to have more information on how dune bryophyte communities would change under a scenario of climatic change (REF). At this stage, it is still uncertain whether the projected climatic alterations will alter the structure and functioning of coastal dune plant communities and further facilitate the spread of invasive alien moss species in these fragile ecosystems. We, therefore, studied coastal dune plant communities along a latitudinal aridity gradient across the western Iberian Atlantic, assessing potential interactions among moss species and between moss cover and vascular plants, and analysed their relationship with abiotic variables to explain the dune bryophyte community and forecast how these communities may respond to climate change.

## 2. Material and Methods

### 2.1 Sampling sites and method

We characterised the vegetation of coastal dunes in 32 sampling sites (SS) along the West Coast of the Iberian Peninsula (Fig. 1), which is characterised by increasing aridity towards the southern end of the gradient. Bryophyte species were identified in 15 sampling sites (out of the 32 SS). Surveys were conducted in medium stabilised dunes, close to the interior of the front dune, where the vegetation is dominated by xerophytic shrubs, from dense/sparse to open structure. Distance to sea of the sampling sites varies from 34 to 360 m due to the geomorphological and structural differences among the studied dune systems. Similar habitat conditions were kept regarding the type of dune system and vegetation community. Sampling was performed during 2017 using the point intercept method (Nunes et al., 2015): at each point, a fine rod was stuck in the ground at a 90° angle and all plant species touching the rod were recorded, by species or by vegetation group (moss, lichen, annual or perennial vascular plant). Cover estimates for individual species or groups were calculated as the proportion of points intercepted per site. In each site, we sampled 4 transects of 50 m parallels to the coastline, every 50 cm (n = 404 points except for SS 7 = 303 and SS 26 = 505) and the soil cover were also recorded (bare soil, lichens, mosses, litter). In addition, one soil sample was collected from top 20 cm in the middle of each transect and merged/bulked between transects to have one composite soil sample for each site.

**Figure 1.**
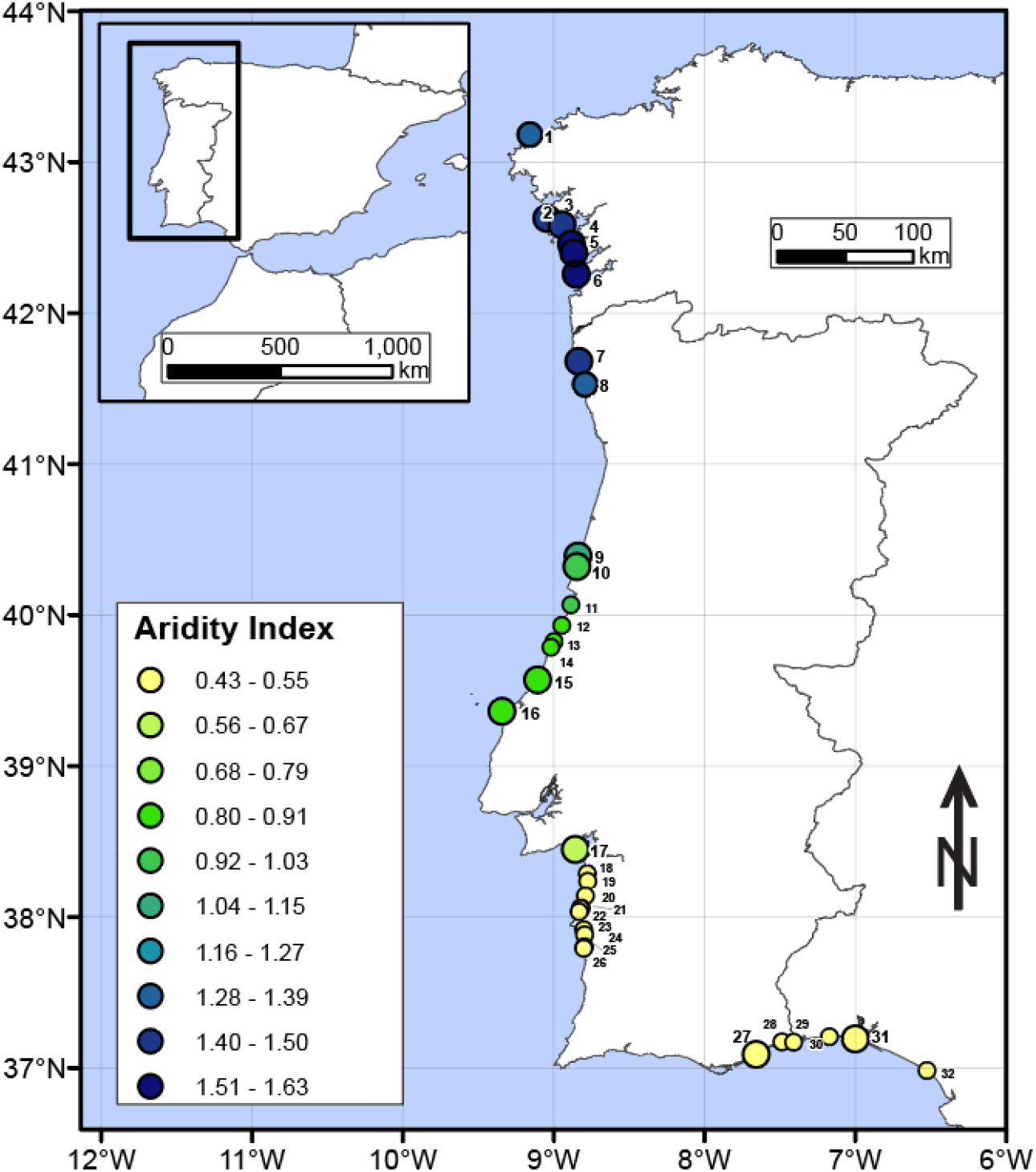
Map showing the study area in the Iberian Peninsula and the location of the 32 sampling sites and the corresponding Aridity Index values (from 0.45 = more arid to 1.63 = less arid) for each Sampling Site according to Fick and Hijmans (2017). The 15 sampling sites where moss species were identified have a larger dot size.

**Figure 2.**
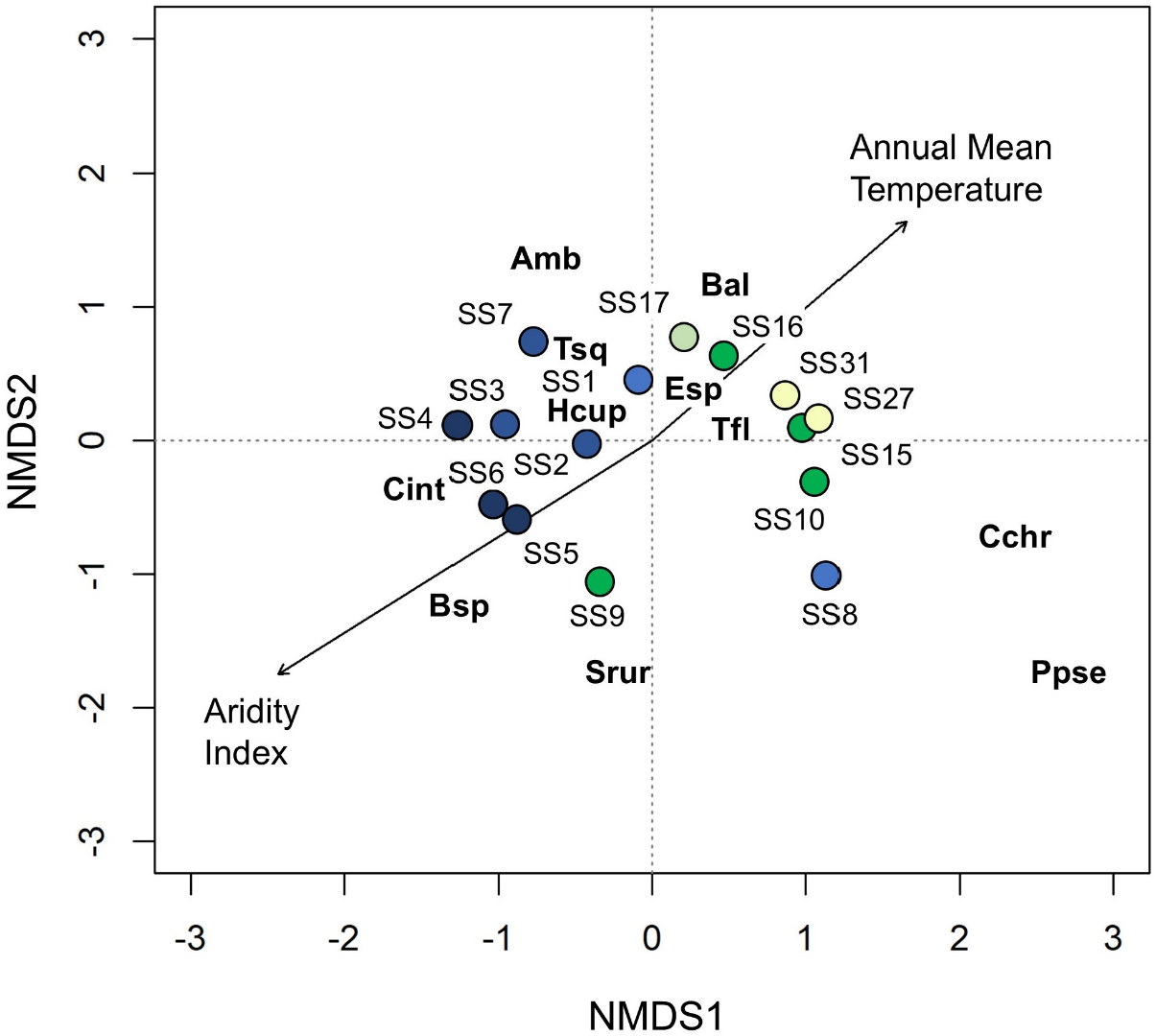
Axes 1 and 2 of the 2-dimensional non-metric multidimensional scaling ordination of sampling sites (SS) based on moss cover (NMDS1 and NMDS2). The final stress was 0.12. Blue, green, pale green and yellow circles are study sampling sites according to their Aridity Index (see Figure 1). Arrows reflect the main gradients identified by the ordination: Aridity Index (r^2^= 0.67 p <0.001) and Annual Mean Temperature (r^2^= 0.40 p <0.05). Species codes: *Amb*.: *Amblystegiaceae; B.a*.: *Brachytecium albicans; B.sp: Bryum sp*.; *C.c*.: *Campyliadelphus chrysophyllus; C.i*.: *Campylopus introflexus; E.sp*.: *Eurhynchium sp; H.c*.: *Hypnum cupressiforme; P.p*.: *Ptychostomum pseudotriquetrum; S.r.:Syntrichia ruralis; T.f*.: *Tortella flavovirens; T.s*.: *Tortella squarrosa*.

### 2.2 Environmental variables

Climate and soil variables were considered to account for the effects of environmental and human disturbances on both moss cover and communities. Data of 19 bioclimatic variables and Aridity Index (AI) were extracted from http://www.worlclim.org/ database (Fick and Hijmans, 2017) from the global aridity database (https://cgiarcsi.community/2019/01/24/global-aridity-index-and-potential-evapotranspiration-climate-database-v2/; Trabucco and Zomer, 2009) adopted by the United Nations, respectively. Soil samples were analysed at Faculdade de Ciências da Universidade de Lisboa (Portugal): pH and soil organic matter content (OM) in the laboratory of Ecology, and total nitrogen (N) and total carbon (C) in SIIAF (Stable Isotopes and Instrumental Analysis Facility).

### 2.3. Data analysis

The main gradients of moss composition were described with a NMDS ordination performed on a matrix of sampling sites by species with the function metaMDS of R Package vegan (Oksanen *et al*. 2013). Only the 15 SS where moss species were identified were included in the ordination analysis. Data were submitted to Wisconsin double standardization (species are first standardized by maxima and then sites by site totals). The Bray and Curtis method was used to measure the distance/similarity between sites. To select the main factor affecting moss composition, we analysed the relationship between the NMDS ordination and the potential explanatory variables through vector fitting (e.g. Chozas et al., 2015). Then, those variables presenting significant correlations were overlaid in the NMDS ordination (McCune & Grace, 2002; Oksanen, 2009). For each set of potential explanatory variables studied (climate and soil) multicollinearity was handled by dropping collinear covariates (Graham 2003; Zuur et al. 2010) when correlated at |Spearman r| > 0.7 (Dormann et al., 2012). Finally, Generalized Additive Models (GAMs) were performed to characterise the relationships of the most important variables conditioning the species composition gradients identified from NMDS analyses with the cover of the mosses using the mgcv software package (Wood 2006).

To determine the relationship between biotic and environmental variables Spearman correlations were performed. In addition, to study vegetation co-occurrence patterns, we used data on the presence/absence at each intercept point of the transects of each vegetation group total cover (i.e. mosses, lichens, perennials and annuals) for the 32 study sites, and also each moss species cover for the 15 sites where the species were identified. Only species or vegetation group data with at least 5% coverage per site were considered. Thus, four incidence matrices were obtained for each sampling site, corresponding to the four transects carried out. These four matrices were joined to configure a single incidence matrix per beach. C-scores were calculated for these incidence matrices to assess co-occurrence patterns (as proposed by Stone & Roberts, 1990). The degree of spatial aggregation of vegetation types and moss species was assessed through the standardized effect size (SES), calculated as indicated in Gotelli & McCabe (2002). SES values greater than 0 indicate spatial segregation among vegetation/species, while smaller values indicate spatial aggregation (Maltez-Mouro et al., 2010). Thus, values above > 2 are considered as competition between groups/species and below < 2 as facilitation. Correlations and their significance were obtained with the cor.test function of the stats package (R Core Team 2018), and the C-score was performed with the cooc_null_model function of the EcoSimR package (Gotelli et al., 2015). All statistical analyses were performed with the statistical program RStudio (RStudio Team 2018).

## 3. Results

The absolute frequencies of mosses, lichens, annuals, perennials, and different moss species at every sampling site are shown in Supplementary Table S1. Mosses were present on 27 out of the 32 sampling sites studied. Total moss cover was highly correlated with the Aridity Index, showing also a significant positive association with climate variables related to precipitation and/or humidity, and a negative one with those related to temperature and/or evapotranspiration (Table 2). For edaphic variables, there is a significant correlation with % N and % C. Regarding biotic factors, total moss coverage only seems to have a significant negative association with plant litter and a positive association with lichen cover.

**Table 2.**
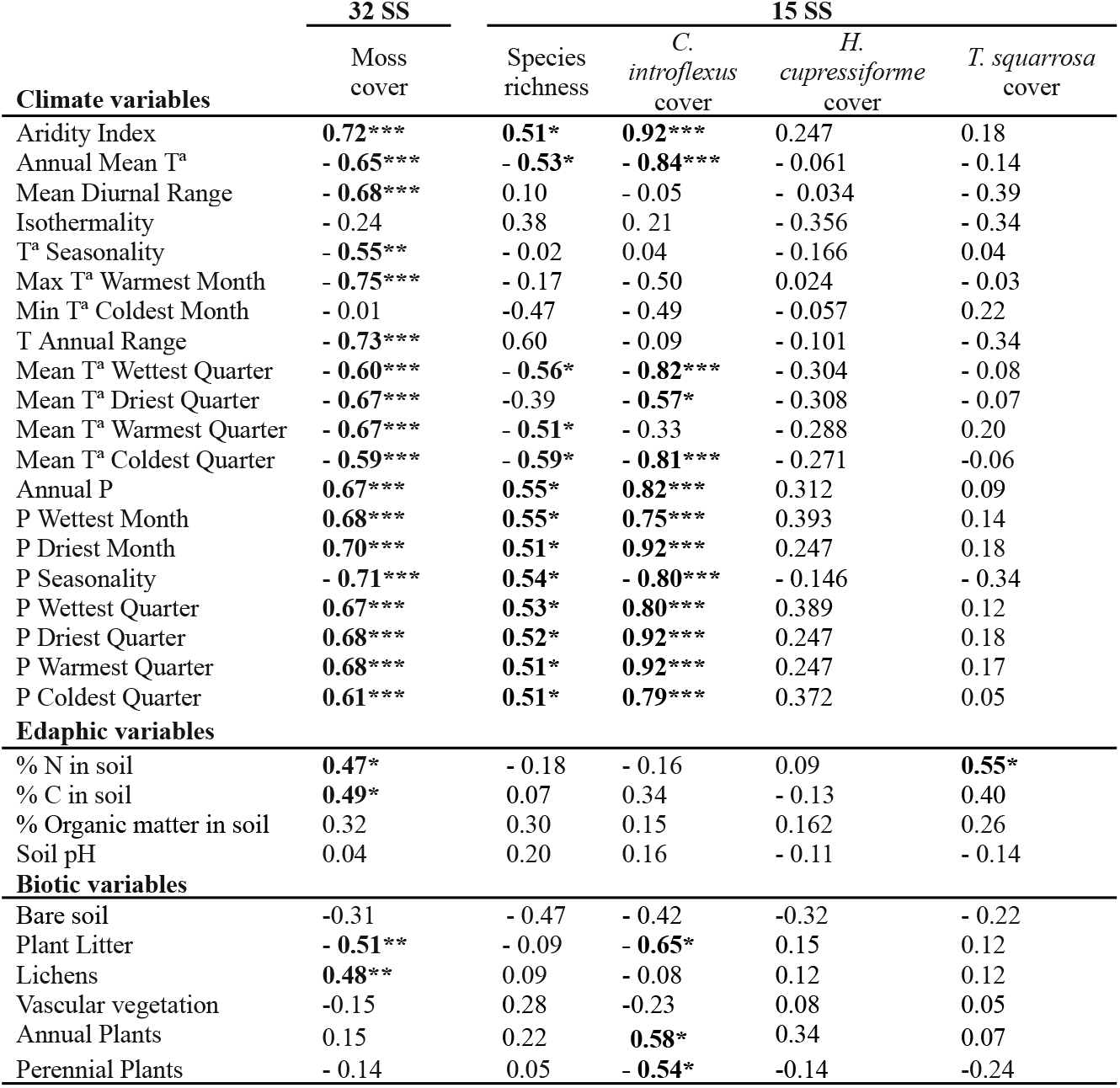
Correlation coefficients (Spearman ρ) between total moss cover, species richness, *Campylopus introflexus, Hypnum cupressiforme and Tortella squarrosa* cover and climate, edaphic and biotic variables. T^a^ = temperature and P = precipitation. Vascular vegetation annuals + perennials. Significances in bold: * p < 0.05; ** p < 0.01; *** p < 0.001.

As shown in Table S1, eleven species of bryophytes were found on the 15 sampling sites where they were characterized. However, most of these species were only present in one or two SS and with very low frequencies. Thus, the correlation analyses have only been performed with *Campylopus introflexus, Hypnum cupressiforme*, and *Tortella squarrosa*. While *T. squarrosa* is present in all sampled sites, *H. cupressiforme* is present in six, and *C. introflexus* is present in 10, only when aridity decreases above AI = 0.68. The cover of the alien invasive moss *C. introflexus* shows a pattern of associations very similar to that shown by total moss coverage (*i.e*. with significant correlations with similar signs for most of the climatic variables). In the case of *T. squarrosa*, its cover is correlated only with the percentage of N in the soil. The association between these two variables is positive, as is the case for total moss cover. Regarding the biotic factors, the cover of *T. squarrosa* seems to have no association with any of them, while the cover of *C. introflexus* seems to have a negative association with the presence of plant litter and perennial plants and positive with annual plant cover.

Regarding the moss community dynamic analyses, the two-dimensional NMDS ordination based on the moss species cover data, with a final stress of 0.12, described the variations in moss cover following an aridity gradient (Figure 3). Correlation analyses confirmed that the main variable describing the gradient among communities was aridity (AI, r^2^= 0.67 p < 0.0001), albeit together with Annual Mean Temperature (AMT, r^2^= 0.40 p < 0.05).

**Figure 3.**
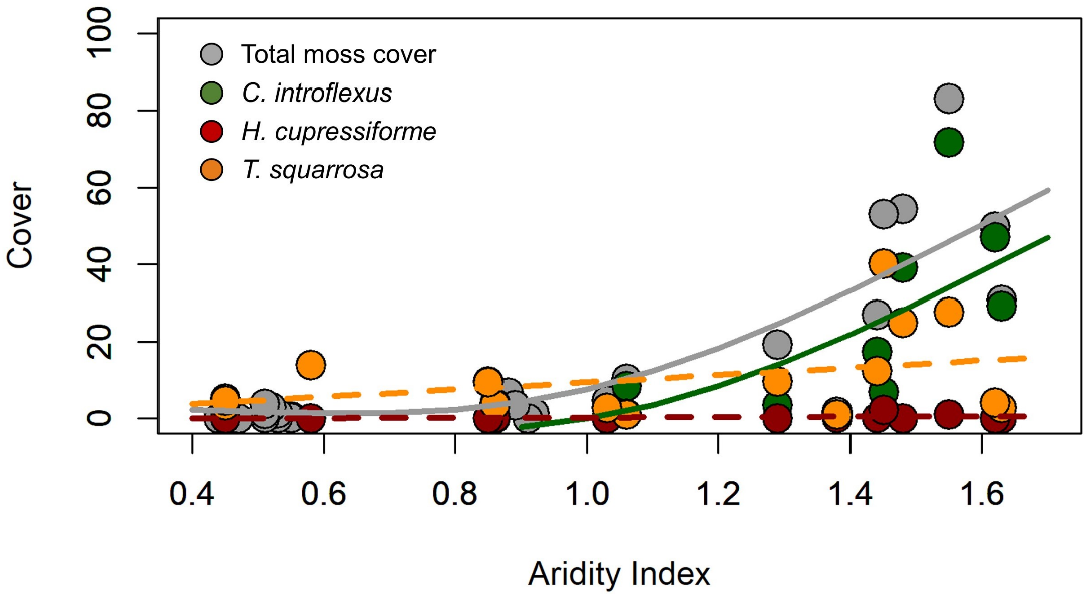
Relationships between the Aridity Index and i) the percentage of total cover of mosses along the 32 study sites (in grey), and ii) the percentage of cover of *C*. introflexus (in dark green*), H. cupressiforme*, (in dark red) and *T. squarrosa* (in orange) in the 15 sampling sites were moss species were identified. Solid lines represent the main trend of a GAM with statistical significance, while dashed lines represent no statistically significant relationships.

The GAMs performed to characterize the relationships between the main variable determining community gradients identified, AI, and the total cover of mosses of the three more abundant species (*C. introflexus, H. cupressiforme*, and *T. squarrosa*), only identified a significant relationship between the aridity and i) the total cover of mosses (Deviance explained = 71.8%, p < 0.001, k = 1.917, n=32) and ii) the cover of *C. introflexus* (De = 60.5%, p < 0.01, k = 1.805, n=15) (Figure 3).

Regarding co-occurrence patterns, there are a few significant (p < 0.05) interactions between mosses and other vegetation (Figure 4). We found facilitation between mosses and lichens in 3/27 SS. On the other hand, it seems that bryophytes compete with perennial plants in 2/27 cases and with annual plants in 1/27. In SS where moss species were identified (n= 15), co-occurrence patterns of the two most abundant moss species show competition between them in 1/15.

**Figure 4.**
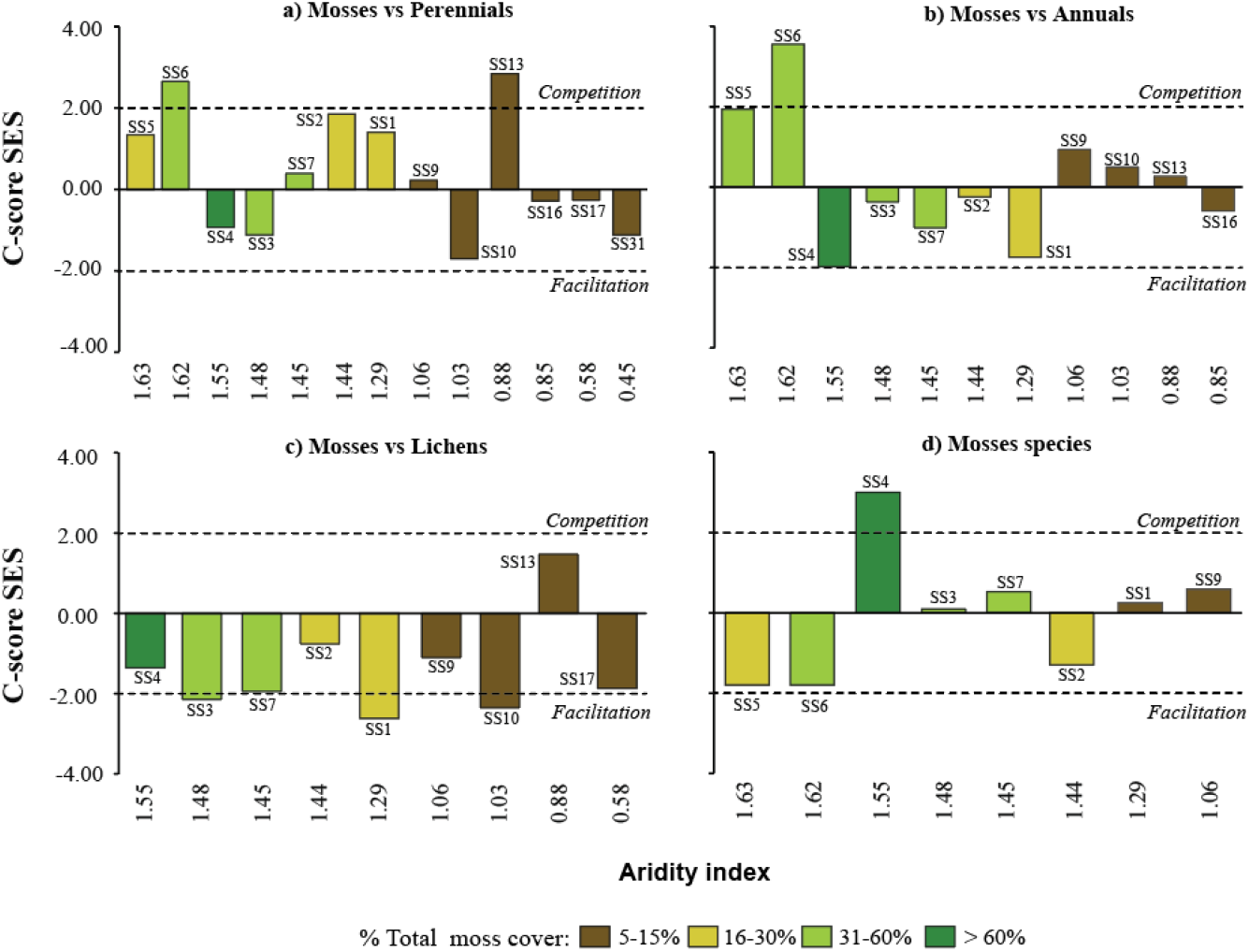
C-score for co-occurrence patterns and standardized effect size (SES) for each vegetation group (i.e., mosses, lichens, perennials, and annuals), and moss species along the aridity gradient with at least 5% cover of each group/species. The aridity index is between 1.63 (less arid) and 0.45 (more arid). The different colours of the columns show the percentage of total moss cover. The dashed lines show the value above which competition between groups/species (> 2) or facilitation (< - 2) is considered to exist.

## 4. Discussion

Our results show that coastal dune moss community dynamics are mainly determined by aridity and temperature along the Western Iberian Atlantic Coast. Although mosses are present in most of the studied sites, their cover and diversity are low and decline towards the most arid and warm areas. These findings differ from the existing literature about European coastal dunes, which reports a high abundance and species richness of mosses (e.g. Jun & Rozé 2004; Callaghan & Ashton, 2007; Provoost et al., 2011). Despite this information, our results are consistent with moss physiology, as their poikilohydric character makes them better adapted to humidity conditions than to drought and heat (Furness & Grime, 1982), showing a lower temperature optimum than higher plants (Glime, 2007). It should be considered that, biogeographically, only the first eight sampling stations are placed in the southernmost part of the Eurosiberian region, while the rest belong to the Mediterranean region (EEA, 2011). In general, these areas tend to be drier than the northern European coasts further where the other studies were conducted. This would lead to higher evaporation rates, shorter periods of photosynthetic activity and more rapid desiccation, which has already been shown to affect moss growth rates and abundance (He et al. 2016). Therefore, this decrease in species number and frequency towards the south (with the subsequent increase in aridity) is expected. Indeed, moss diversity peaks northwards compared with other plant groups (Mateo et al., 2016; Ronquillo et al. 2023).

An alternative explanation for these results could be the uncontrolled urbanisation that has taken place in the coastal areas of southern Europe since the beginning of the century, leading to the loss of vegetation and the disappearance of 70% of the European coastal dune systems (Brown and McLachlan, 2010; Gómez-Pina et al., 2002). However, we did not find any relationship between land-use or perturbation and the cover and diversity of mosses in our study area (data not shown) but it may be worth bearing in mind that the recovery time of bryophytes in disturbed dune environments can vary significantly depending on levels of aridity (Kammann et al., 2022; Ladrón de Guevara and Maestre, 2022).

These dynamics in the bryophyte community could be even more extreme under climate change conditions (Mendoza-González et a., 2013). Nonetheless, according to the literature, biotic interactions with other plants could help avoid losing bryophyte diversity and abundance in the context of a changing climate and stressful environment (e.g. Ingerpuu et al., 2005; Brooker et al., 2008). The Stress Gradient Hypothesis (SGH) suggests that facilitation occurs under high-stress conditions and competition under low-stress conditions (Bertness and Callaway, 1994), while Doxford et al. (2013) found that under these stress conditions, biotic interactions may also depend on population growth rate. However, our data can only partially be explained by SGH (Bertness and Callaway, 1994) and based on total cover (not growth rate), competition between mosses and annuals was only found in one of the eleven SS, located at one end of the aridity gradient (SS6, Fig. 4) and with intermediate-high vegetation cover (30-60%). Competition between mosses and perennials also occurs in this SS6, as well as in SS13, located in the middle of the gradient, with low plant cover (5-15%, Fig. 4). On the other hand, moss and lichen cover show a positive association, as is often observed due to their role as colonising organisms in the early stages of primary succession in dune systems (e.g Jun & Rozé, 2004). Therefore, facilitation interactions between them would be expected to be found independently of the aridity gradient. Even so, we only found facilitation between mosses and lichens in 3 of the 9 SS (Fig. 4): in SS1 and SS3, the northernmost study sites, with the rather low aridity values and intermediate-high cover (i.e. 30-60% and > 60% respectively); and SS10, with intermediate aridity values and low cover (between 5-15%).

This lack of evidence for interactions may be due to the inconsistent variability in the presence/absence of vegetation at the sampled stations (Table 1). Vaz et al. (2020), already reported the lack of relationship between the diversity patterns of bryophytes, lichens and vascular plants in dunes with contrasting coastal dynamics, regardless of biogeographic context or anthropogenic pressures. Furthermore, the joint effects of regional and local environmental gradients may be buffering or masking these biotic processes that may be occurring at finer scales as outlined by Brooker et al. (2008) and Vaz et al. (2015). Consequently, in these few interactions found, all the hypotheses proposed by previous studies on the existence of competition/facilitation both at the extremes and at intermediate values of the aridity gradient, as well as those dependent on total vegetation cover, are fulfilled, with no common pattern along the gradient. Given the limited bryophyte-plant vascular interactions, it is unlikely that these interactions will help mitigate potential bryophytes losses caused by climate change events in the future.

As regards the increase of alien species aggravated by climate change (Cogoni et al. 2011), the invasive moss *Campylopus introflexus* only appears when the AI increased and was not found in the 5 SS of maximum aridity (Table 1, Fig. 3). This is consistent with its significantly positive correlations with rainfall, and negative correlations with temperature variables (Table 2). This indicates that *C. introflexus* is likely unable to compete effectively with perennial species, while it can coexist with herbaceous annuals due to a lower competition for light and nutrients (Corbin & D’Antonio, 2004). This dynamic is especially pronounced in scenarios where perennial species are present in high densities and annuals in low densities, as found in our study sites (e.g., Coomes et al. 2002). All this confirms that *C. introflexus*, as an invasive species, prefers open areas or is related to more disturbed ecosystems, with less vegetation and high anthropogenic influence (see Sérgio et al., 2018). Out of the 8 SS where co-occurrence between *T. squarrosa* (very common in coastal dunes) and *C. introflexus* was analysed, only in SS4 was competition between them found (Fig. 4). This SS has the highest cover of the two species and is at the extreme minimum aridity (Table 1 and Fig. 3), so, given these results, we confirm that SGH is met but conditional on optimal conditions for both species. In SS where the number of occurrences of one or both species is low, it is likely that the coverage required for competitive interactions is not achieved.

## 5. Conclusions

The dynamics of the dune moss community along the western Iberian Atlantic coast are mainly determined by aridity and temperature while biotic interactions do not seem to drive these dynamics. In the current context of climate change and increasing aridity, these moss communities will decrease their total coverage in dune environments. Given their minor importance, their interactions with other plant groups are unlikely to compensate for this loss. Under this scenario, it is also reasonable to assume that the optimal conditions for the different moss species will shift towards higher latitudes due to the lower rate of aridity and, therefore, the situations of competition between moss species will be increasingly displaced to the north. On the other hand, it is possible that the invasion of *Campylopus introflexus*, which seems to be associated with lower aridity rates, will slow down in this region. Even so, more detailed studies will be needed to evaluate the alterations in the structural and functional characteristics of these fragile coastal habitats of Community conservation concern.

## Acknowledgements

The authors would like to thank J.R. Aboal for his help with Figure 1. This work was supported by the projects UNITED (CGL2016-78070-P, funded by Spanish AEI and FEDER, UE) and by the project ChangeTracker, FCT-PTDC/AAG-GLO/0045/2014. ZV contributed to this study with a postdoctoral research grant awarded by the Xunta de Galicia (Spain; Modalidade B-2019) but currently is supported by the María Zambrano Programme of the Spanish Ministry of Universities. SC gratefully acknowledges the support of the Fundação para a Ciência e Tecnologia (FCT), through the strategic project UIDB/00329/2020 - https://doi.org/10.54499/UIDB/00329/2020 granted to cE3c-Centre for Ecology, Evolution and Environmental Changes, Faculdade de Ciências, Universidade de Lisboa. SC was supported by the project BIOSOLAR through a postdoctoral fellowship. SLD and JM were supported by PhD Grants FCT-PD/BD/114438/2016 and SFRH/BD/131924/2017 respectively. P. Giráldez is grateful to the Spanish Ministerio de Ciencia, Innovación y Universidades for a grant awarded within the Programa de Formacion de Profesorado Universitario (FPU 2018 [grant number FPU18/04134]). JH was funded by Project NICED, grant PID2022-140985NB-C21 funded by MCIN/AEI/ 10.13039/501100011033 / FEDER, EU.

---

**Table 1SM.** Absolute frequencies of mosses, lichens, annual and perennial plants, and the different moss species on the sampled sites. n= 404 except in B7= 303 and B26= 505 (marked with asterisk). Moss species: *Amb*.: *Amblystegiaceae; B.a*.: *Brachytecium albicans; B.sp: Bryum sp*.; *C.c*.: *Campyliadelphus chrysophyllus; C.i*.: *Campylopus introflexus; E.sp*.: *Eurhynchium sp; H.c*.: *Hypnum cupressiforme; P.p*.: *Ptychostomum pseudotriquetrum; S.r.:Syntrichia ruralis; T.f*.: *Tortella flavovirens; T.s*.: *Tortella squarrosa*.

| SS | Absolute frequency |  |  |  | Absolute frequency of each moss species |  |  |  |  |  |  |  |  |  |  |
| --- | --- | --- | --- | --- | --- | --- | --- | --- | --- | --- | --- | --- | --- | --- | --- |
|  | Moss | Lichen | Annual | Perennia<br>l | *Amb.* | *B. a.* | *B. sp* | *C. c.* | *C. i.* | *E. sp* | *H. c.* | *P. p* | *S. r.* | *T. f.* | *T. s.* |
| SS1 | 78 | 15 | 28 | 361 | 0 | 23 | 0 | 0 | 14 | 4 | 0 | 0 | 0 | 8 | 39 |
| SS2 | 109 | 14 | 3 | 191 | 0 | 0 | 0 | 0 | 70 | 0 | 1 | 0 | 0 | 0 | 50 |
| SS3 | 220 | 51 | 27 | 207 | 0 | 0 | 0 | 0 | 158 | 0 | 0 | 0 | 0 | 0 | 101 |
| SS4 | 336 | 9 | 38 | 233 | 0 | 0 | 0 | 0 | 290 | 0 | 4 | 0 | 0 | 0 | 112 |
| SS5 | 124 | 0 | 23 | 227 | 0 | 0 | 1 | 0 | 118 | 0 | 0 | 0 | 1 | 0 | 11 |
| SS6 | 201 | 0 | 58 | 166 | 0 | 0 | 0 | 0 | 190 | 0 | 0 | 0 | 0 | 0 | 17 |
| SS7* | 161 | 25 | 43 | 148 | 22 | 0 | 0 | 0 | 21 | 0 | 7 | 0 | 0 | 0 | 122 |
| SS8 | 7 | 0 | 79 | 297 | 0 | 0 | 0 | 0 | 3 | 0 | 1 | 1 | 0 | 0 | 4 |
| SS9 | 42 | 115 | 7 | 237 | 0 | 0 | 0 | 2 | 34 | 0 | 5 | 0 | 2 | 2 | 4 |
| SS10 | 19 | 53 | 5 | 244 | 0 | 0 | 0 | 10 | 1 | 0 | 0 | 0 | 0 | 0 | 11 |
| SS11 | 6 | 6 | 4 | 234 | - | - | - | - | - | - | - | - | - | - | - |
| SS12 | 0 | 1 | 4 | 310 | - | - | - | - | - | - | - | - | - | - | - |
| SS13 | 27 | 89 | 1 | 356 | - | - | - | - | - | - | - | - | - | - | - |
| SS14 | 14 | 2 | 1 | 398 | - | - | - | - | - | - | - | - | - | - | - |
| SS15 | 16 | 4 | 15 | 251 | 0 | 0 | 0 | 0 | 0 | 0 | 1 | 0 | 0 | 0 | 15 |
| SS16 | 39 | 0 | 6 | 228 | 0 | 0 | 0 | 0 | 0 | 0 | 0 | 0 | 0 | 0 | 38 |
| SS17 | 56 | 65 | 0 | 359 | 0 | 0 | 0 | 0 | 0 | 0 | 0 | 0 | 0 | 0 | 56 |
| SS18 | 2 | 0 | 1 | 267 | - | - | - | - | - | - | - | - | - | - | - |
| SS19 | 1 | 19 | 5 | 274 | - | - | - | - | - | - | - | - | - | - | - |
| SS20 | 1 | 0 | 0 | 273 | - | - | - | - | - | - | - | - | - | - | - |
| SS21 | 1 | 0 | 1 | 242 | - | - | - | - | - | - | - | - | - | - | - |
| SS22 | 6 | 7 | 7 | 327 | - | - | - | - | - | - | - | - | - | - | - |
| SS23 | 12 | 2 | 20 | 293 | - | - | - | - | - | - | - | - | - | - | - |
| SS24 | 1 | 0 | 10 | 353 | - | - | - | - | - | - | - | - | - | - | - |
| SS25 | 5 | 0 | 11 | 376 | - | - | - | - | - | - | - | - | - | - | - |
| SS26* | 19 | 0 | 6 | 431 | - | - | - | - | - | - | - | - | - | - | - |
| SS27 | 14 | 36 | 0 | 224 | 0 | 0 | 0 | 0 | 0 | 0 | 0 | 0 | 0 | 0 | 14 |
| SS28 | 0 | 0 | 12 | 212 | - | - | - | - | - | - | - | - | - | - | - |
| SS29 | 0 | 0 | 232 | 144 | - | - | - | - | - | - | - | - | - | - | - |
| SS30 | 0 | 0 | 51 | 214 | - | - | - | - | - | - | - | - | - | - | - |
| SS31 | 21 | 1 | 4 | 356 | 2 | 0 | 0 | 0 | 0 | 0 | 0 | 0 | 0 | 0 | 19 |
| SS32 | 0 | 0 | 26 | 133 | - | - | - | - | - | - | - | - | - | - | - |

## Notes

### Competing Interest Statement

The authors have declared no competing interest.

